# Rendering protein mutation movies with MutAmore

**DOI:** 10.1101/2023.09.15.557870

**Authors:** Konstantin Weissenow, Burkhard Rost

## Abstract

The success of *AlphaFold2* in reliable protein three-dimensional (3D) structure prediction, assists the move of structural biology toward studies of protein dynamics and mutational impact on structure and function. This transition needs tools that qualitatively assess alternative 3D conformations. We introduce *MutAmore*, a bioinformatics tool that renders individual images of protein 3D structures for, e.g., sequence mutations into a visually intuitive movie format. *MutAmore* streamlines a pipeline casting single amino-acid variations (SAVs) into a dynamic 3D mutation movie providing a qualitative perspective on the mutational landscape of a protein. By default, the tool first generates all possible variants of the sequence reachable through SAVs (L*19 for proteins with L residues). Next, it predicts the structural conformation for all L*19 variants using state-of-the-art models. Finally, it visualizes the mutation matrix and produces a color-coded 3D animation. Alternatively, users can input other types of variants, e.g., from experimental structures. *MutAmore* samples alternative protein configurations to study the dynamical space accessible from SAVs in the post-AlphaFold2 era of structural biology. As the field shifts towards the exploration of alternative conformations of proteins, *MutAmore* aids in the understanding of the structural impact of mutations by providing a flexible pipeline for the generation of protein mutation movies using current and future structure prediction models.

## Introduction

### AI has changed structural biology and its impact

The remarkable success of *AlphaFold2* [1] in effectively predicting protein three-dimensional (3D) structure from sequence is shifting paradigms in structural biology and beyond. *AlphaFold2* combines advanced Artificial Intelligence (AI) with evolutionary information from multiple sequence alignments (MSAs). Reliable 3D predictions for over 200 million proteins [2] have begun to change molecular biology. Concurrently, protein Language Models (pLMs) have emerged as a new approach to represent protein sequences [3-5]. Downstream prediction methods based on pLM embeddings (vectors representing the last hidden layers of the pLMs: Fig. 1 [4]) find new solutions, e.g., in structure prediction methods such as *ESMFold* [6]. Embeddings allow predictions at unprecedented speed [7, 8], often outperforming state-of-the-art methods [9] using the combination of evolutionary information introduced three decades ago for secondary structure prediction [10] and eclipsed by *AlphaFold2*.

**Fig. 1:**
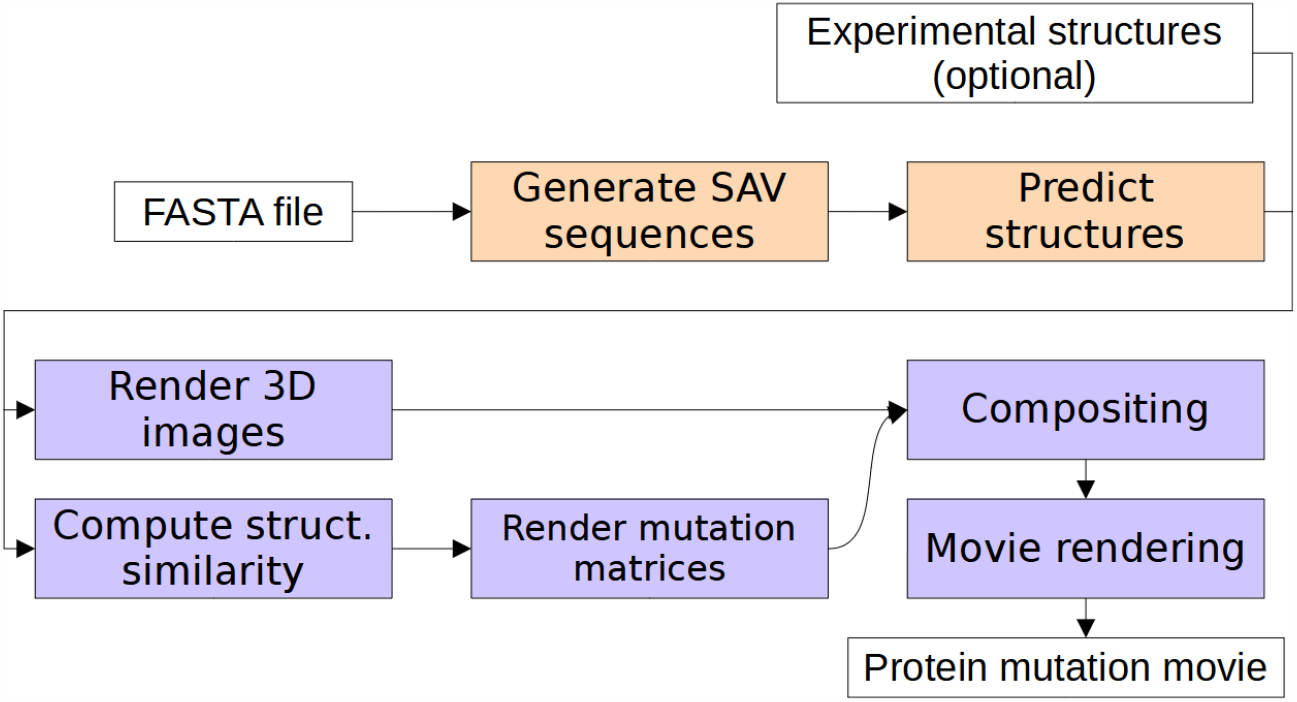
MutAmore pipeline. The tool generates mutated (all possible SAVs) versions of the input sequences and predicts 3D structure for each, e.g., using ColabFold [33] or ESMFold [6]. Optionally, experimental structures for mutants can be input as an alternative to predictions. *MutAmore* computes the structural difference between each mutant and the predicted wild-type structure and renders mutation profiles along with 3D visualizations. After merging both, the tool renders the final protein mutation movie (PMM). *MutAmore* can be run in a two-step process, e.g., predicting structure (orange) on a server machine and running the rendering steps (blue) on a desktop machine.

### From available 3D model to dynamics?

In the post-*AlphaFold2* era, structural biology moves toward a deeper exploration of alternative protein conformations and mutational landscapes [11]. Many proteins significantly change function upon minor sequence changes [12-14]. Understanding these changes will be critical to unlocking the complexities of protein function, protein evolution, and disease progression at the molecular level.

Despite its immense success, *AlphaFold2* often fails to correctly predict the effect of missense mutations upon 3D structure [7, 8, 15]. Even if it correctly captured such effects, it would still be too resource-intensive to be used to explore the mutational space of an average protein, e.g., by evaluating all possible single amino-acid substitutions (SAVs) even for short proteins [7, 8]. Structure predictions based on pLMs such as ESMFold or EMBER2 [7] offer both the necessary speed and promise higher sensitivity to small changes in input sequences (Fig. 2 [8]).

**Fig. 2:**
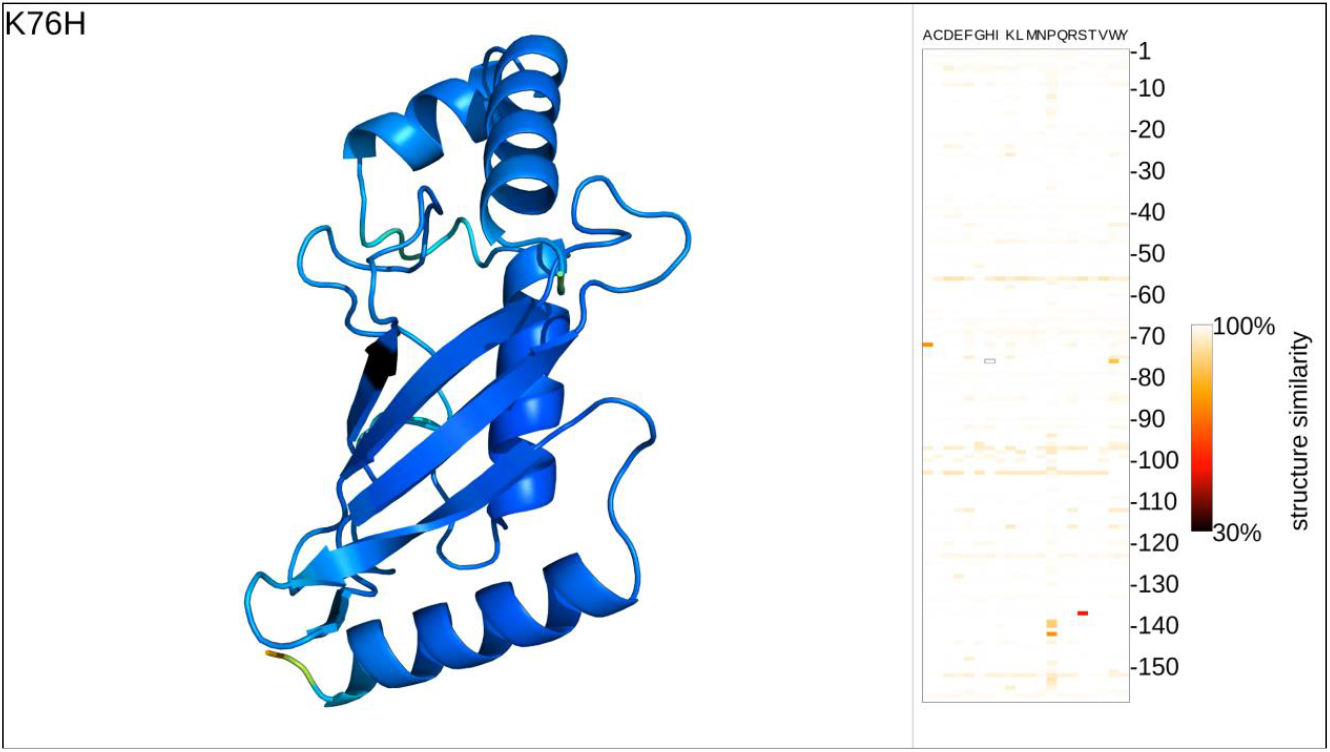
Single frame of protein mutation movie (PMM). *<u>Left panel:</u>* 3D visualization of the prediction for one single amino-acid variant (SAV). The respective SAV is indicated in the top-left corner (residue position 83: native lysin (K) mutated into isoleucine (I)) and the affected residue is rendered in black in the visualization. All other residues are colored by the *AlphaFold2* predicted confidence in the *AlphaFold2* color scheme (blue: high confidence; yellow-red: low confidence). *<u>Right panel:</u>* the mutation profile shows the structural difference between mutant and wild-type structure with residue indices running along the vertical axis and substitution amino-acids on the horizontal. The currently displayed SAV in each frame is indicated by a black border. The movie frame shown was rendered in a screen resolution of 3840x2160.

Many methods have been developed to predict the effects of sequence variants upon protein function, including SIFT [16], PolyPhen [17], SNAP2 [18], GEMME [19], DeepSequence [20], Packpred [21], Tranception [22] and VESPA [23] (for a longer list: [22, 24]). On the other hand, tools such as I-Mutant [25, 26], FoldX [27], PoPMuSiC [28], DUET [29], or INPS-MD [30] aim at predicting the structural impact of mutations by providing estimates for changes in stability, folding or dynamics, which greatly helps our understanding of disease emergence. Despite convincing examples for how to use the numerical output from such methods to rationalize on static images about possible dynamical changes [21, 29, 31], none of these methods directly displays the actual structural change in the 3D conformation.

Imagine we had fast and accurate structure predictions for all L*19 SAV mutants of a protein with L residues. How to visualize those data? Today, no comprehensive and accessible visualization tools for alternative protein conformations are available. To fill this void, we introduced *MutAmore* (**MutA**tion **mo**vie **re**nderer), a tool that provides a pipeline rendering a mutated protein sequence into a dynamic 3D protein mutation movie (PMM), thereby making the analysis of mutational landscapes accessible and tangible. The implementation and results presented highlight the potential of *MutAmore* to fill a growing need within the structural biology field. Its efficient visualizations could aid research of protein dynamics, function, evolution, and disease mechanisms.

## Implementation

*MutAmore* is designed to create an animated visualization of the mutational landscape of proteins. Inputting a protein amino acid sequence (or a set thereof) as a FASTA file [32], *MutAmore* first generates all possible single amino acid variants (SAVs) for this sequence(s) and uses a structure prediction model to predict 3D structure for each SAV (Fig. 1). We provide a ready-made interface to *ESMFold* [6] and *ColabFold/AlphaFold2* [33] along with documentation on how to easily use any other prediction model inputting FASTA files and outputting PDB files. Optionally, a user can provide experimental structures for some of the mutants to *MutAmore*. These variants are then skipped during the prediction stage.

*MutAmore* computes a mutation profile by assessing the structural divergences between each mutated protein and the (predicted) wild-type structure. The *local Distance Difference Test* (lDDT) [34] tallies scores across all residues to derive a single score for each structure pair (typically wild-type/native vs. SAV). Structures similar to the wild-type have scores near one while divergent structures approach zero. These data are then visualized as a mutation matrix (19 x protein length) using Python-Pillow [35].

3D visualizations of all variants are created via the PyMOL API [36] and color-coded to indicate predicted confidence levels (blue for high confidence, yellow-red for low confidence; following the *AlphaFold2* standard). All recent 3D structure prediction methods include predicted lDDT values as confidence scores, including systems which do not build on top of the AlphaFold2 architecture, such as RoseTTAFold [37] and EMBER3D [8]. PyMOL aligns all 19*L SAV structures to the original wild-type prediction prior to rendering to maintain uniformity in viewing angles in the final animated visualization.

*MutAmore* then assembles the final frames of the animation in residue index order (from first at the N-terminus to the last at the C-terminus), merging the 3D renderings with the mutation profile to create the PMM with ffmpeg [38]. Rendered at a rate of 19 frames per second, each residue remains in focus for one second and shows all potential SAVs for this position in the protein. The mutated position in each frame is rendered in black in both the 3D visualization and the mutation profile. Details of the SAVs displayed in each frame are indicated in the top-left corner through the standard single-letter amino acid code: XnY meaning that the wild-type amino acid X at residue position n is mutated to amino acid Y (Fig. 2: K76H).

When users provide experimental structures – or otherwise their own labels – in a PDB-formatted file, the visual output highlights the differences between predictions and experimental models by showing the latter at full opacity while applying slight transparency to the former, giving the visual impression of “filling in the gaps” between known structures with predictions.

Users can adapt the resolution of the final animation to their needs, e.g., choosing high-quality for publications or lower resolution clips for web sharing. The mutation profile automatically scales to the specified vertical resolution for optimal visual interpretability. The 3D visualization rendered by PyMOL automatically chooses a zoom level, which allows enough space to accommodate structural changes caused by mutations. *MutAmore* lets experienced users override the zoom level manually where needed.

*MutAmore* also lets users render subsets of the most impactful mutations, e.g., top-50: those with the highest effect upon 3D. In this mode, the framerate is slightly reduced (slowed down) for better visual comparison.

Many advanced structure prediction systems require robust and substantial GPU resources. Therefore, *MutAmore* provides an option for a two-step process: computation of all mutation predictions using a server machine, and subsequent processing and rendering of the animated visualization on a desktop computer.

## Results

We evaluated *MutAmore* on an Intel Xeon Gold 6248 CPU with a NVidia Quadro RTX 8000 GPU (48GB) using twelve proteins ranging in lengths from 72 to 639 residues, including both globular and membrane proteins. Although *AlphaFold2* [1] or its faster spin-off *ColabFold* [33] outperform *ESMFold* [6], the latter seems slightly better at capturing the effects of SAVs upon structure [7, 8]. We also used ESMFold to predict 3D structures because it is substantially faster which mattered for the 55,879 SAVs in all twelve samples. Then we generated movies at the default resolution of 1280x720 (720p) and 3840x2160 (4K, Table 1).

**Table 1:** Benchmark of *MutAmore* in 720p and 4K resolution for twelve proteins. *.

|  | Runtime in seconds |  |
| --- | --- | --- |
|  | 720p resolution | 4K resolution |
| *Prediction (ESMFold)* | 1,423,814 |  |
| *3D rendering* | 26,893 | 106,829 |
| *Structural similarity* | 4,008 | 4,224 |
| *Mutation profile rendering* | 4,472 | 16,871 |
| *Compositing final frames* | 4,730 | 33,032 |
| *Movie rendering* | 172 | 1,012 |
| *Total (rendering)* | 40,275 | 161,968 |
| *Total (rendering + prediction)* | 1,464,089 | 1,585,782 |

\* Runtime of individual pipeline steps in seconds for screen resolutions 720p and 4K, summed up over twelve proteins with lengths between 72 and 639 residues (total number of SAVs/3D predictions: 55,879; total number of seconds playtime: 2,941). Structure prediction with ESMFold [6] and structural similarity computation using lDDT [34] are naturally unaffected by larger image sizes, but 3D rendering with PyMOL [36], compositing (putting together the individual image pieces) and movie rendering (encoding individual frames to a movie) scale with a factor closer to the increase in screen estate. Tables S1-S12 in the Supporting Online Materials break down the runtime for each of the twelve proteins while Fig. S13 visualizes the scaling of runtime with protein length. Note: the 1.4 million seconds required for the *ESMFold* 3D predictions amounted to about 16 days.

The time required for structure predictions substantially varies with the method used. ColabFold, utilized for the five smallest proteins only, accumulated over a week of GPU time, given the non-linear scale with protein length, this rapidly becomes infeasible for longer proteins. ESMFold obtained predictions for all twelve proteins in about 16 days, with the bulk of the computation time dedicated to the longest samples (11 days for the protein with 639 residues). In contrast, the predictions for the nine smaller proteins were computed by ESMFold in 30 hours. The tremendous increase in runtime by protein length (Fig. S13) is due to GPU memory limitations. Structure prediction systems such as ESMFold compute multiple samples in parallel, to fill memory as efficiently as possible. For shorter proteins, this allows batches of up to dozens of simultaneous predictions. Longer sequences require more computation time and more memory. This limits the number of samples that can be processed concurrently, leading to an exponential increase in total runtime. Overall, these numbers highlight the need for future structure predictions with both increased speed and memory efficiency to properly explore the mutational landscape of longer proteins.

Given the 3D predictions, creating the animated visualizations is considerably faster. The majority of *MutAmore’s* processing time is devoted to the rendering of 3D visualizations, followed by the composition of final frames.

After generating structure predictions on our server hardware, we applied *MutAmore’s* rendering pipeline on a consumer grade laptop with an Intel Core i7 6700HQ CPU to compare performance with the server environment. Performance decreased by roughly 30%, showing that *MutAmore* fits to readily available hardware, at least for shorter proteins.

Enhancing the resolution required additional runtime. Rendering at 4K increased the processing time for the 3D visualization, compositing, and rendering by a factor of three to seven over rendering at 720p (Table 1), but generally less than the increase in amount of pixels (4K/720p=9x). The time for computing structural similarity and for generating the profile did not differ much between 720p and 4K (Table 1). We provide a detailed breakdown of the runtime of all pipeline steps on the twelve individual samples in the Supporting Online Materials (SOM: Tables S1-S12). Even at 4K, the total processing time for *MutAmore* remained substantially below that needed to predict structures even with *ESMFold* for all but the shortest protein sequences (Fig. S13).

## Limitations

A profound consequence of all attempts to visualize the dynamics of 3D objects lies in the obstruction of internal parts. For instance, buried residues and changes of local regions around these will remain obscured from our 3D movies. For some proteins, such as beta barrels, a cleverly chosen viewing angle might provide a better perspective, but globular proteins do not provide such an alternative. Transparency might address such issues, but transparency tends to cause a lack of depth perception and too much visual clutter to clearly comprehend the visual information being presented. Thus, it remains unclear how to show internal structural changes of proteins in a visually concise manner apart from going back to two-dimensional distance maps, which are only intuitive to a well-trained structural expert.

Another implicit limitation for using 3D predictions for all 19-non-native SAVs is in the substantial demand on computing resources of today’s prediction methods. This becomes particularly challenging for long proteins (Fig. S13), *MutAmore* would greatly benefit from future prediction systems that are leaner without having to sacrifice performance [8].

Too few high-resolution experimental structures establish the effect of point mutations upon 3D to evaluate how well prediction methods capture SAV effects. Proxying deep mutational scanning data [39] might suffice to establish correlations between observed and predicted impact upon function without probing how well methods predict the effect of SAVs upon 3D structure and dynamics. Once more experimental data of variant structures will be available, *MutAmore* could be extended to serve as a benchmarking tool for the sensitivity of structure prediction.

A webserver for *MutAmore* might ease the access for users with less experience in computational biology. While we are looking for the resources to realize such a project, we provide a Google Colab Notebook linked on the *MutAmore* website and allows the creation of PMM’s without the need of a local installation.

## Conclusions

We designed *MutAmore* to bridge a crucial gap in the post-AlphaFold era. By rendering conceivable and visually comprehensible protein mutation movies (PMMs) of single amino-acid substitutions (SAVs), *MutAmore* enhances the exploration and understanding of alternative protein conformations brought on by mutations. This is particularly significant in translating the depth and complexity of the protein mutational landscape. Our benchmark demonstrated the efficiency and versatility of *MutAmore* even for high-definition 4K video settings. The tool effectively balances computational load by allowing multi-step operation across multiple systems, ensuring usability across varied system capabilities. We hope that structural biology increasingly shifts toward routine analysis of alternate protein conformations. *MutAmore* supports such a more dynamic perspective on protein structures and might aid analyzing protein function and studying protein evolution.

## Abbreviations used

3D: three-dimensional (coordinates)
3D structure: three-dimensional coordinates of protein structure
AI: Artificial Intelligence
API: application programing interface
embeddings: fixed-size vectors derived from pre-trained pLMs
GPU: graphical processing unit
PDB: Protein Data Bank
PIDE: percentage pairwise sequence identity
pLM: protein Language Model
PMM: protein mutation movie

## Availability

*MutAmore* is publicly available at https://github.com/kWeissenow/MutAmore.

## Acknowledgements

Thanks to Tim Karl (TUM) for invaluable help with hardware and software; to Nikita Kugut (TUM) for support with many other aspects of this work; to Michael Heinzinger (TUM) for helpful discussions and comments on work and manuscript. Furthermore, we thank all those who make experimental and predicted structures and other resources publicly available, in particular, thanks to DeepMind (AlphaFold2) and Meta (ESMFold), as well as to the creators and community of PyMOL and ffmpeg.

