## Supplemental files for "Rendering protein mutation movies with MutAmore"

### Supporting Online Material (SOM) for:
#### Rendering protein mutation movies with MutAmore

Konstantin Weissenow[1,2] & Burkhard Rost [1,3]

1 TUM (Technical University of Munich) Department of Informatics, Bioinformatics & Computational Biology - i12, Boltzmannstr. 3, 85748 Garching/Munich, Germany
2 TUM Graduate School, Center of Doctoral Studies in Informatics and its Applications (CeDoSIA), Boltzmannstr. 11, 85748 Garching, Germany
3 Institute for Advanced Study (TUM-IAS), Lichtenbergstr. 2a, 85748 Garching/Munich, Germany & TUM School of Life Sciences Weihenstephan (WZW), Alte Akademie 8, Freising, Germany

### Table of Contents for SOM

### Short description of SOM

In figures S1-S12 we show screenshots of all protein mutation movies used for benchmarking. Corresponding tables S1-S12 indicate runtimes for all individual samples, including prediction time using ESMFold (Lin et al., 2022) and all *MutAmore* rendering pipeline steps.

Figure S13 visualizes the scaling of runtime for prediction and rendering steps with sequence length.

### Material

#### Fig. S1: Protein mutation movie screenshot: Calmodulin-1:

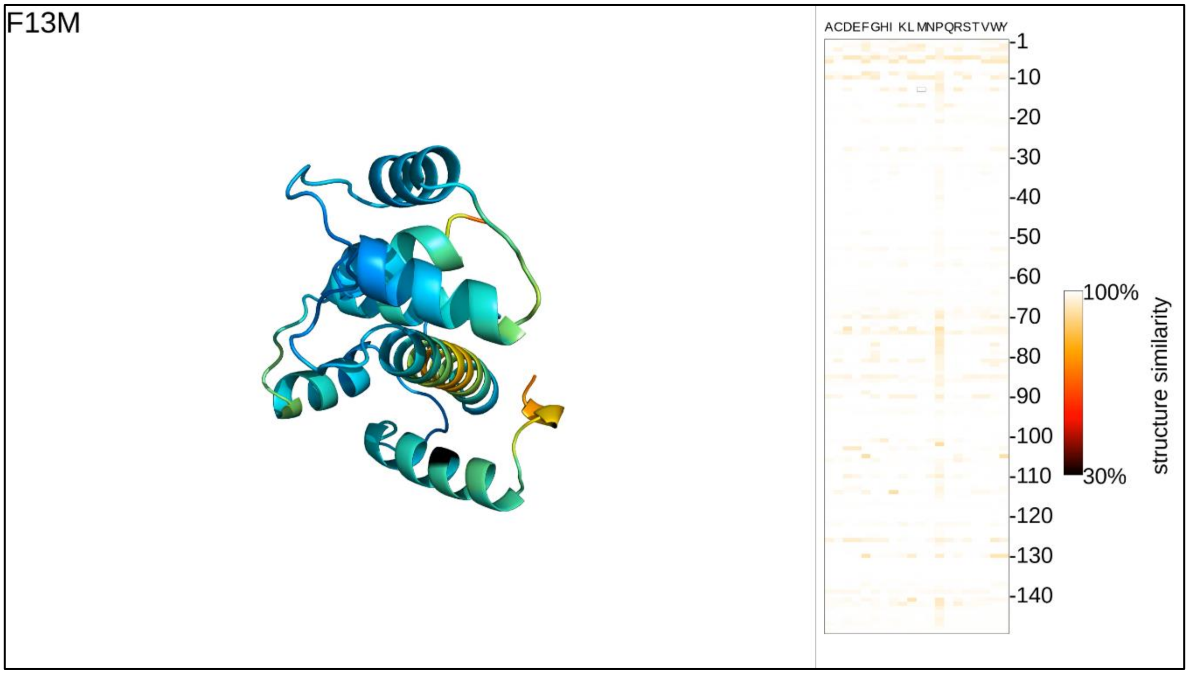

**Fig. S1: Protein mutation movie screenshot: Calmodulin-1.** Single frame of a protein mutation movie for Calmodulin-1.

#### Table S1: Runtimes for Calmodulin-1

| | Runtime (in sec) | |
|---|---|---|
| | 720p resolution | 4K resolution |
| *Prediction (ESMFold)* | 5,346 | |
| *3D rendering* | 957 | 3,389 |
| *Structural similarity* | 116 | 116 |
| *Mutation profile rendering* | 187 | 483 |
| *Compositing final frames* | 217 | 1,350 |
| *Movie rendering* | 9 | 51 |
| *Total (rendering)* | 1,486 | 5,389 |
| *Total (all)* | 6,832 | 10,735 |

**Fig. S2: Protein mutation movie screenshot: Translation initiation factor IF-1:**

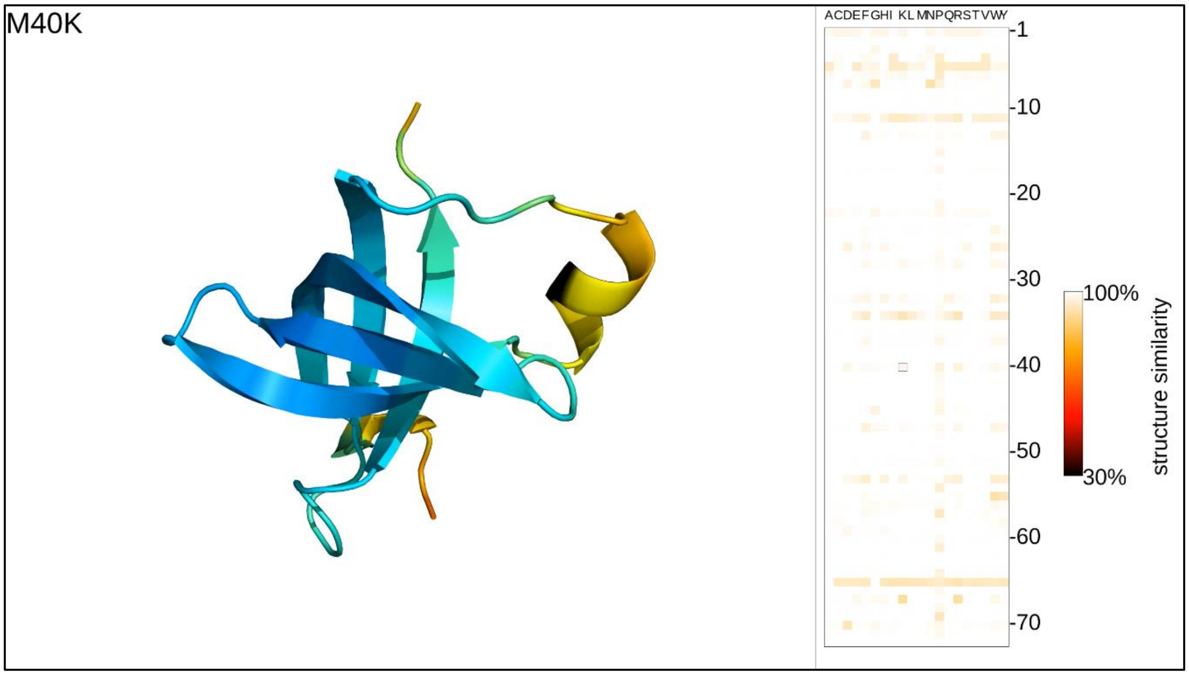

**Fig. S2: Protein mutation movie screenshot: Translation initiation factor IF-1.** Single frame of a protein mutation movie for Translation initiation factor IF-1.

**Table S2: Runtimes for Translation initiation factor IF-1**

| | Runtime (in sec) | |
|---|---|---|
| | 720p resolution | 4K resolution |
| *Prediction (ESMFold)* | 591 | |
| *3D rendering* | 357 | 1,387 |
| *Structural similarity* | 33 | 34 |
| *Mutation profile rendering* | 63 | 177 |
| *Compositing final frames* | 97 | 598 |
| *Movie rendering* | 5 | 26 |
| *Total (rendering)* | 555 | 2,222 |
| *Total (all)* | 1,146 | 2,813 |

**Fig. S3: Protein mutation movie screenshot: Translation initiation factor Hras:**

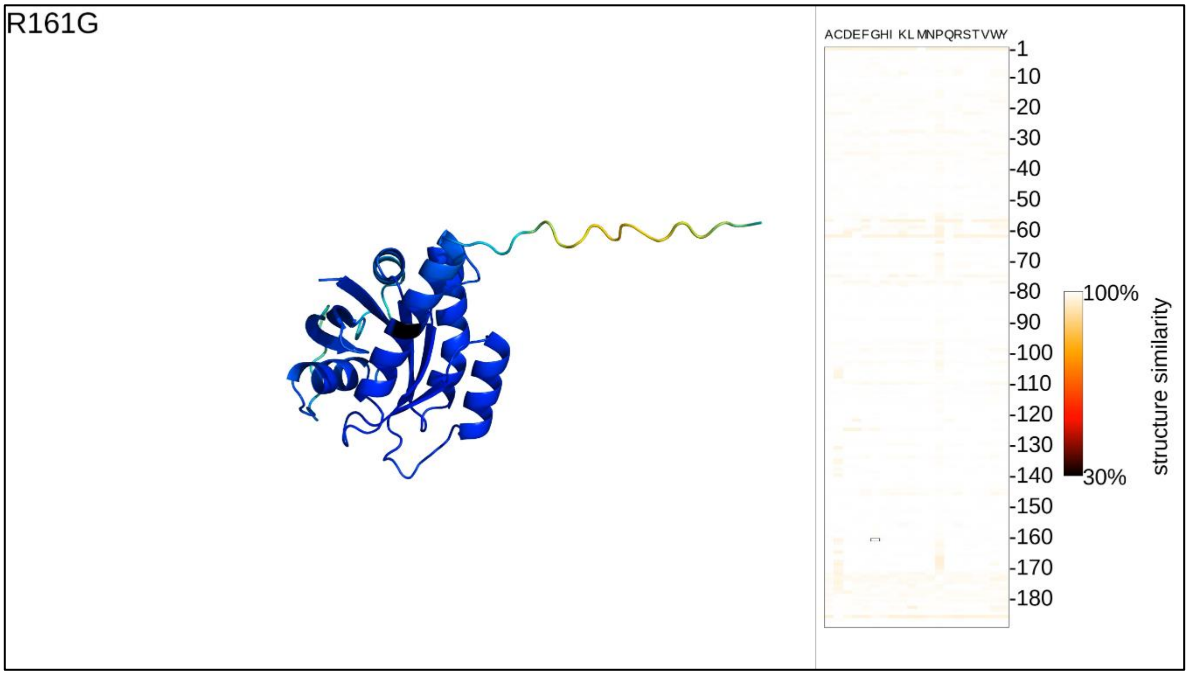

**Fig. S3: Protein mutation movie screenshot: Hras.** Single frame of a protein mutation movie for Hras.

**Table S3: Runtimes for Hras**

| | Runtime (in sec) | |
|---|---|---|
| | 720p resolution | 4K resolution |
| *Prediction (ESMFold)* | 11,198 | |
| *3D rendering* | 1,034 | 3,558 |
| *Structural similarity* | 179 | 177 |
| *Mutation profile rendering* | 272 | 704 |
| *Compositing final frames* | 250 | 1,581 |
| *Movie rendering* | 10 | 65 |
| *Total (rendering)* | 1,745 | 6,085 |
| *Total (all)* | 12,943 | 17,283 |

**Fig. S4: Protein mutation movie screenshot: Yeast ubiquitin:**

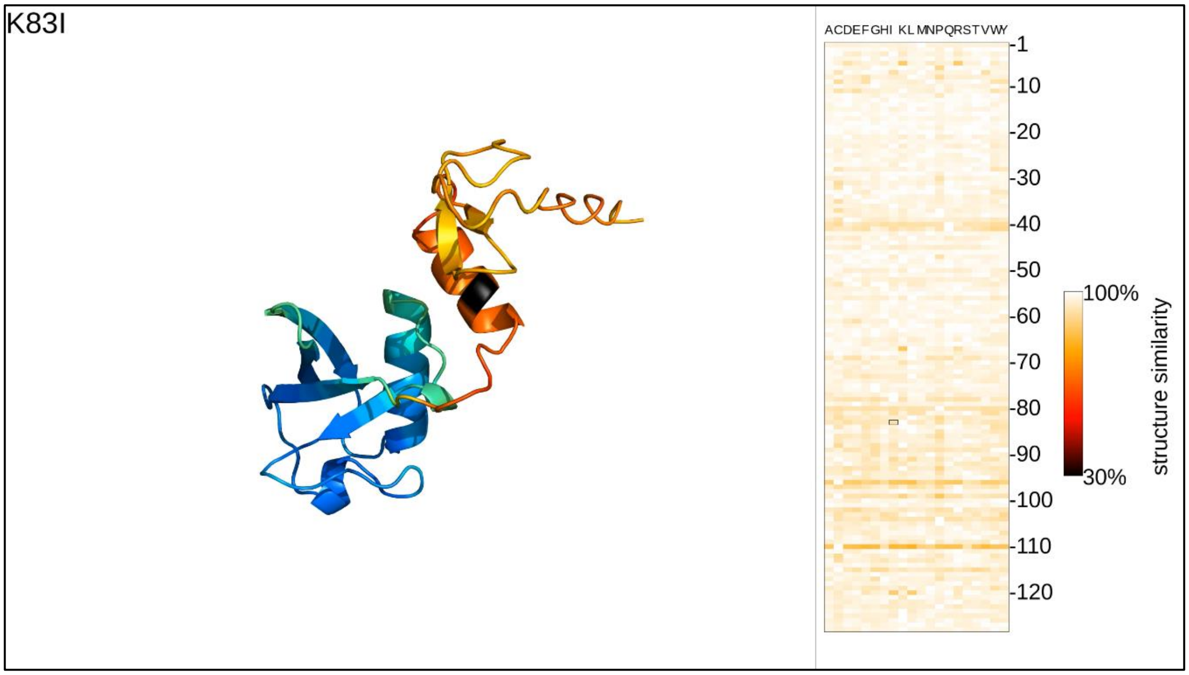

**Fig. S4: Protein mutation movie screenshot: Yeast ubiquitin.** Single frame of a protein mutation movie for yeast ubiquitin.

**Table S4: Runtimes for yeast ubiquitin**

| | Runtime (in sec) | |
|---|---|---|
| | 720p resolution | 4K resolution |
| *Prediction (ESMFold)* | 3,226 | |
| *3D rendering* | 709 | 2,517 |
| *Structural similarity* | 72 | 88 |
| *Mutation profile rendering* | 147 | 391 |
| *Compositing final frames* | 153 | 1,039 |
| *Movie rendering* | 8 | 49 |
| *Total (rendering)* | 1,089 | 4,084 |
| *Total (all)* | 4,315 | 7,310 |

**Fig. S5: Protein mutation movie screenshot: Small ubiquitin-related modifier 1:**

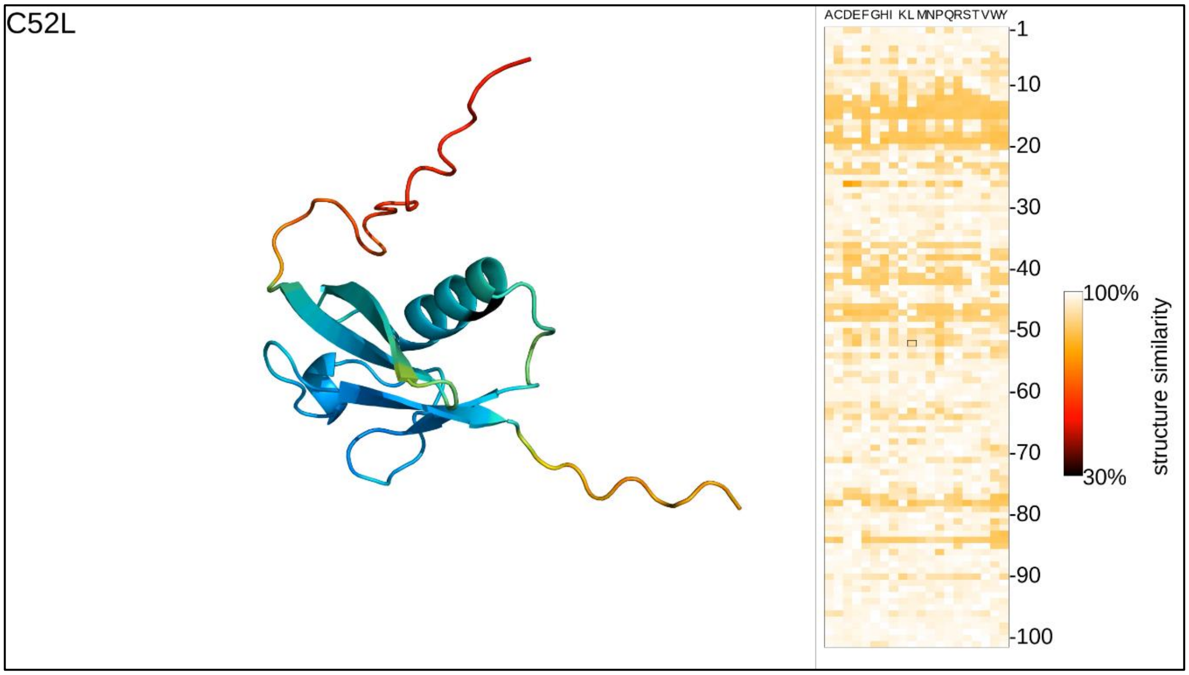

**Fig. S5: Protein mutation movie screenshot: Small ubiquitin-related modifier 1.** Single frame of a protein mutation movie for Small ubiquitin-related modifier 1.

**Table S5: Runtimes for small ubiquitin-related modifier 1**

| | Runtime (in sec) | |
|---|---|---|
| | 720p resolution | 4K resolution |
| *Prediction (ESMFold)* | 1,597 | |
| *3D rendering* | 484 | 1,834 |
| *Structural similarity* | 60 | 59 |
| *Mutation profile rendering* | 99 | 288 |
| *Compositing final frames* | 122 | 815 |
| *Movie rendering* | 6 | 37 |
| *Total (rendering)* | 771 | 3,033 |
| *Total (all)* | 2,368 | 4,630 |

#### Fig. S6: Protein mutation movie screenshot: TIM barrel:

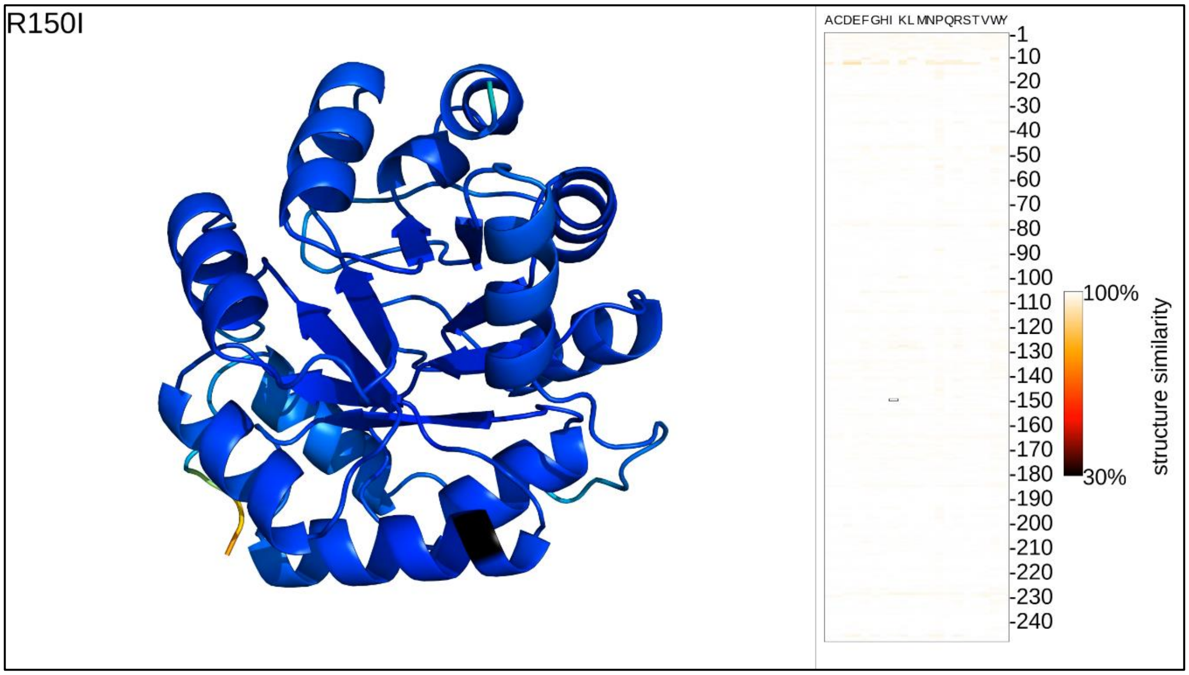

**Fig. S6: Protein mutation movie screenshot: TIM barrel.** Single frame of a protein mutation movie for TIM barrel.

#### Table S6: Runtimes for TIM barrel

| | Runtime (in sec) | |
|---|---|---|
| | 720p resolution | 4K resolution |
| *Prediction (ESMFold)* | 28,261 | |
| *3D rendering* | 2,285 | 9,048 |
| *Structural similarity* | 276 | 273 |
| *Mutation profile rendering* | 396 | 1,093 |
| *Compositing final frames* | 464 | 2,995 |
| *Movie rendering* | 15 | 86 |
| *Total (rendering)* | 3,436 | 13,495 |
| *Total (all)* | 31,697 | 41,756 |

**Fig. S7: Protein mutation movie screenshot: Thiamin pyrophosphokinase 1:**

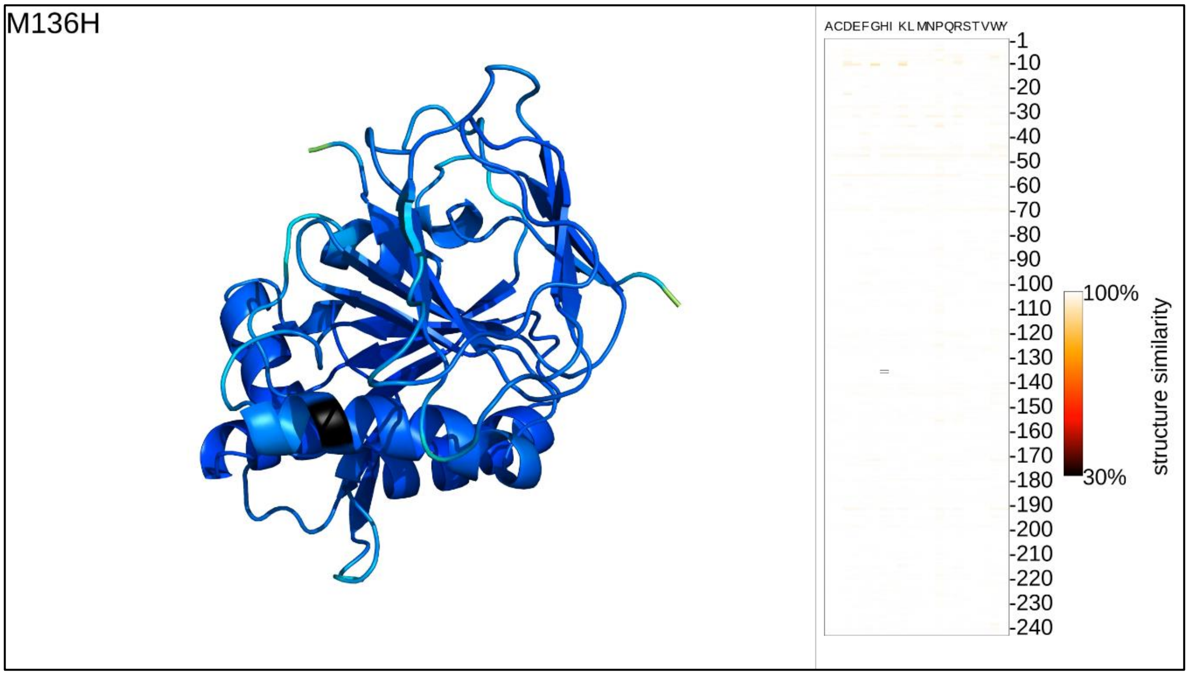

**Fig. S7: Protein mutation movie screenshot: Thiamin pyrophosphokinase 1.** Single frame of a protein mutation movie for Thiamin pyrophosphokinase 1.

**Table S7: Runtimes for Thiamin pyrophosphokinase 1**

| | Runtime (in sec) | |
|---|---|---|
| | 720p resolution | 4K resolution |
| *Prediction (ESMFold)* | 26,481 | |
| *3D rendering* | 2,042 | 6,969 |
| *Structural similarity* | 271 | 273 |
| *Mutation profile rendering* | 398 | 1,075 |
| *Compositing final frames* | 387 | 2,603 |
| *Movie rendering* | 14 | 80 |
| *Total (rendering)* | 3,112 | 11,000 |
| *Total (all)* | 29,593 | 37,481 |

**Fig. S8: Protein mutation movie screenshot: Thiopurine S-methyltransferase:**

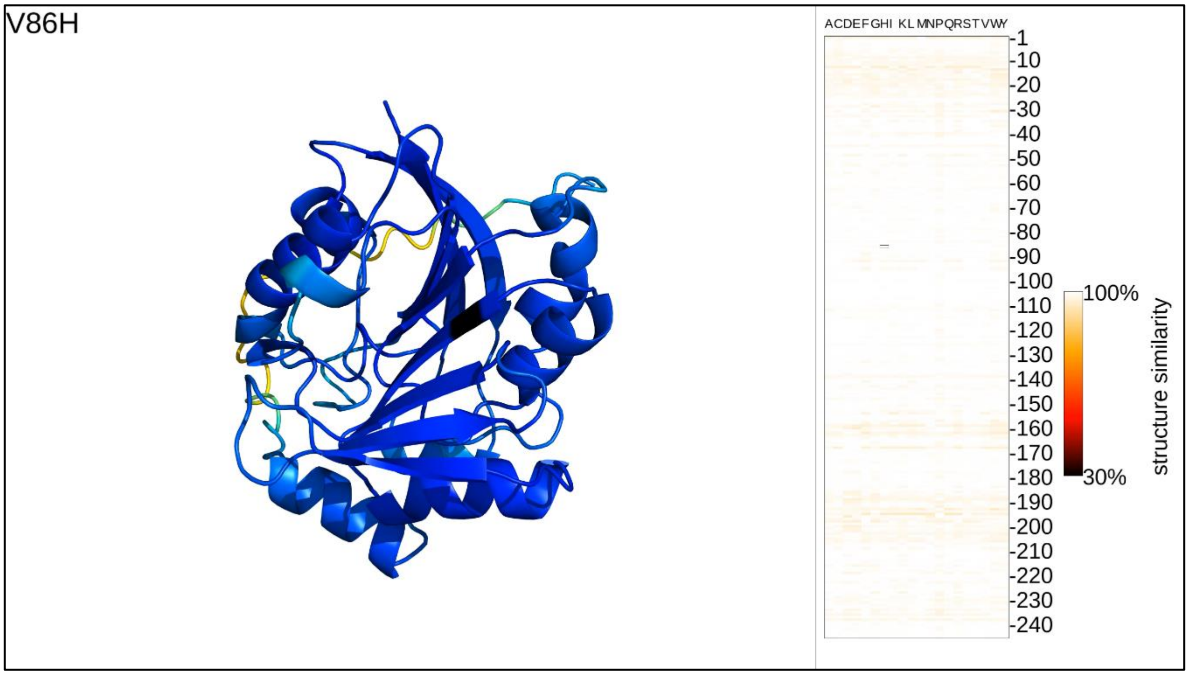

**Fig. S8: Protein mutation movie screenshot: Thiopurine S-methyltransferase.** Single frame of a protein mutation movie for Thiopurine S-methyltransferase.

**Table S8: Runtimes for Thiopurine S-methyltransferase**

| | Runtime (in sec) | |
|---|---|---|
| | 720p resolution | 4K resolution |
| *Prediction (ESMFold)* | 27,296 | |
| *3D rendering* | 1,986 | 6,656 |
| *Structural similarity* | 284 | 286 |
| *Mutation profile rendering* | 402 | 1,086 |
| *Compositing final frames* | 384 | 2,494 |
| *Movie rendering* | 15 | 83 |
| *Total (rendering)* | 3,071 | 10,605 |
| *Total (all)* | 30,367 | 37,901 |

**Fig. S9: Protein mutation movie screenshot: SUMO-conjugating enzyme UBC9:**

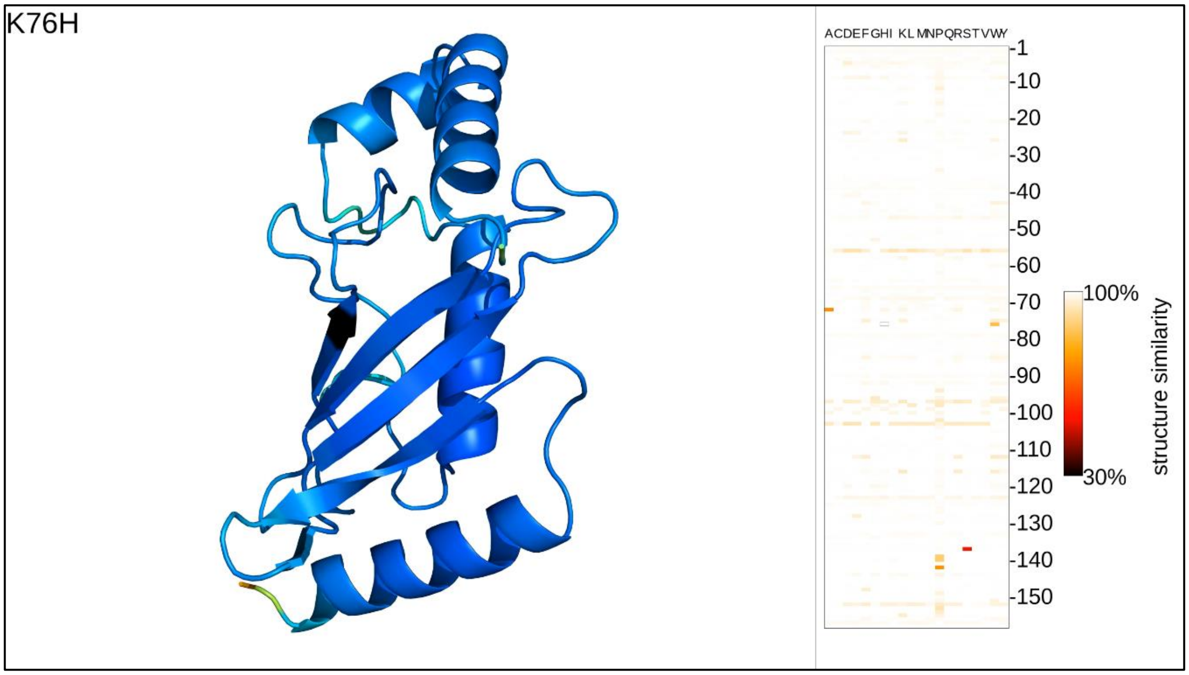

**Fig. S9: Protein mutation movie screenshot: SUMO-conjugating enzyme UBC9.** Single frame of a protein mutation movie for SUMO-conjugating enzyme UBC9.

**Table S9: Runtimes for SUMO-conjugating enzyme UBC9**

| | Runtime (in sec) | |
|---|---|---|
| | 720p resolution | 4K resolution |
| *Prediction (ESMFold)* | 6,338 | |
| *3D rendering* | 1,071 | 4,018 |
| *Structural similarity* | 121 | 134 |
| *Mutation profile rendering* | 197 | 533 |
| *Compositing final frames* | 235 | 1,675 |
| *Movie rendering* | 10 | 55 |
| *Total (rendering)* | 1,634 | 6,415 |
| *Total (all)* | 7,972 | 12,753 |

**Fig. S10: Protein mutation movie screenshot: Xylose transporter XylE from E.coli:**

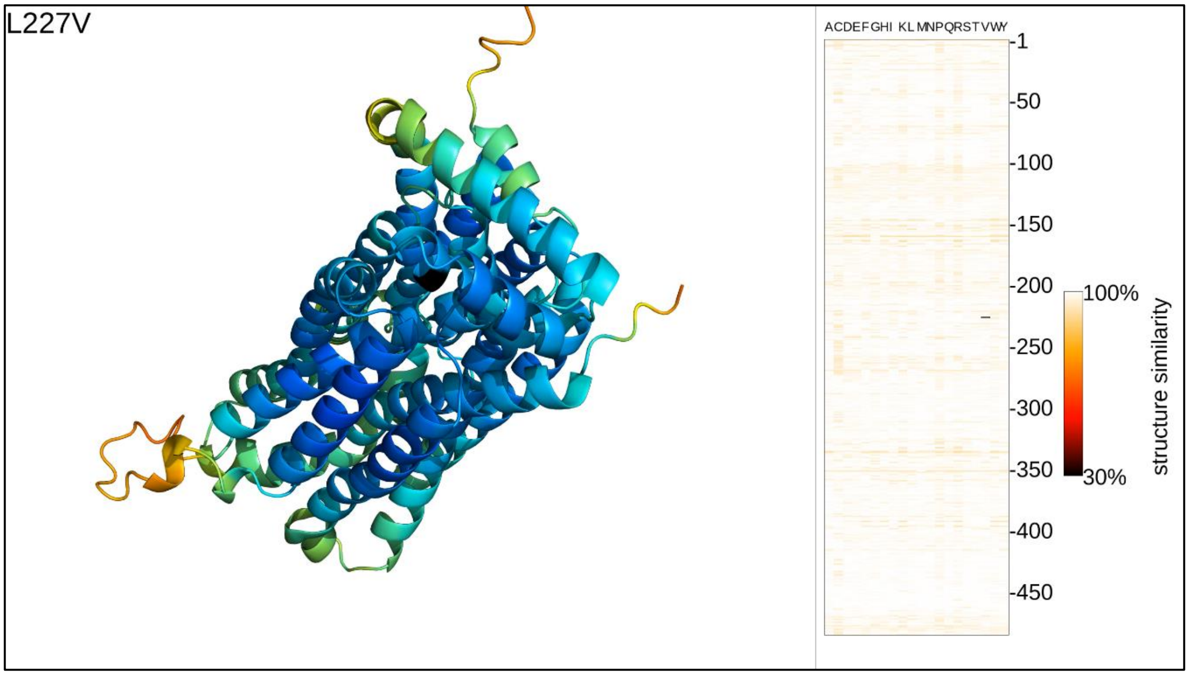

**Fig. S10: Protein mutation movie screenshot: Xylose transporter XylE from E.coli.** Single frame of a protein mutation movie for the xylose transporter XylE from E.coli.

**Table S10: Runtimes for xylose transporter XylE from E.coli**

| | Runtime (in sec) | |
|---|---|---|
| | 720p resolution | 4K resolution |
| *Prediction (ESMFold)* | 321,716 | |
| *3D rendering* | 5,895 | 25,471 |
| *Structural similarity* | 839 | 976 |
| *Mutation profile rendering* | 683 | 4,267 |
| *Compositing final frames* | 853 | 7,208 |
| *Movie rendering* | 30 | 168 |
| *Total (rendering)* | 8,300 | 38,090 |
| *Total (all)* | 330,016 | 359,806 |

**Fig. S11: Protein mutation movie screenshot: E.coli autotransporter beta-domain EspP:**

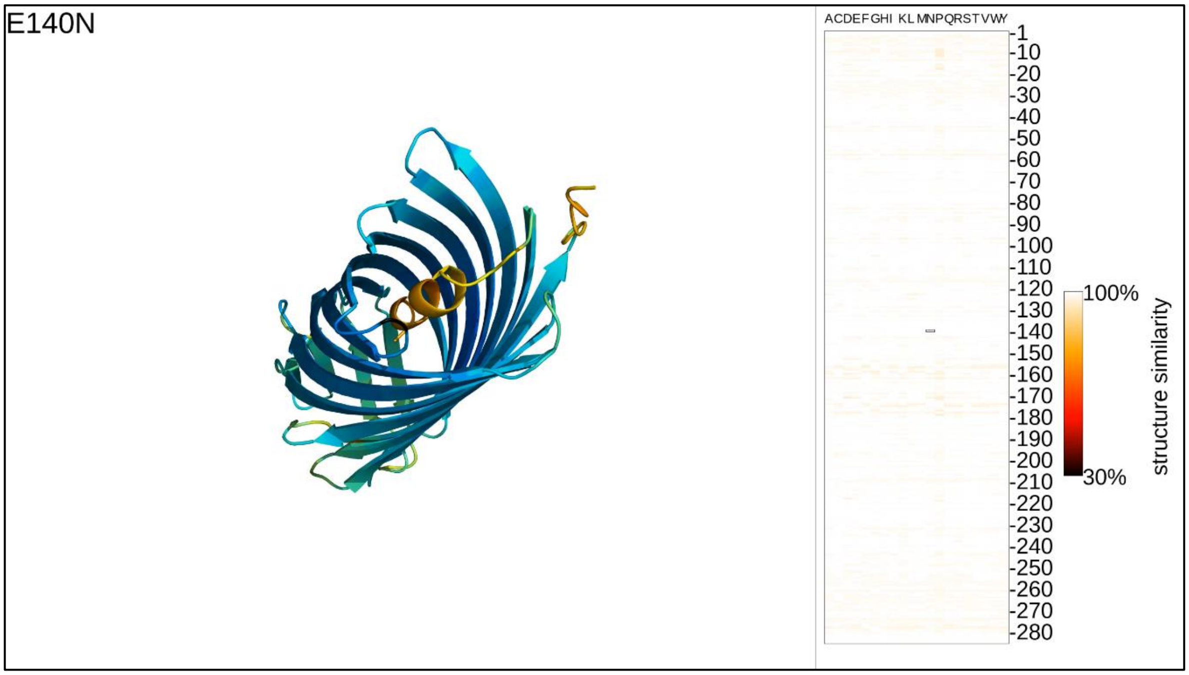

**Fig. S11: Protein mutation movie screenshot: E.coli autotransporter beta-domain EspP.** Single frame of a protein mutation movie for E.coli autotransporter beta-domain EspP.

**Table S11: Runtimes for E.coli autotransporter beta-domain EspP**

| | Runtime (in sec) | |
|---|---|---|
| | 720p resolution | 4K resolution |
| *Prediction (ESMFold)* | 47,429 | |
| *3D rendering* | 2,070 | 8,659 |
| *Structural similarity* | 345 | 386 |
| *Mutation profile rendering* | 499 | 1,557 |
| *Compositing final frames* | 385 | 2,943 |
| *Movie rendering* | 15 | 110 |
| *Total (rendering)* | 3,314 | 13,655 |
| *Total (all)* | 50,743 | 61,084 |

#### Fig. S12: Protein mutation movie screenshot: Colicin I receptor Cir from E.coli:

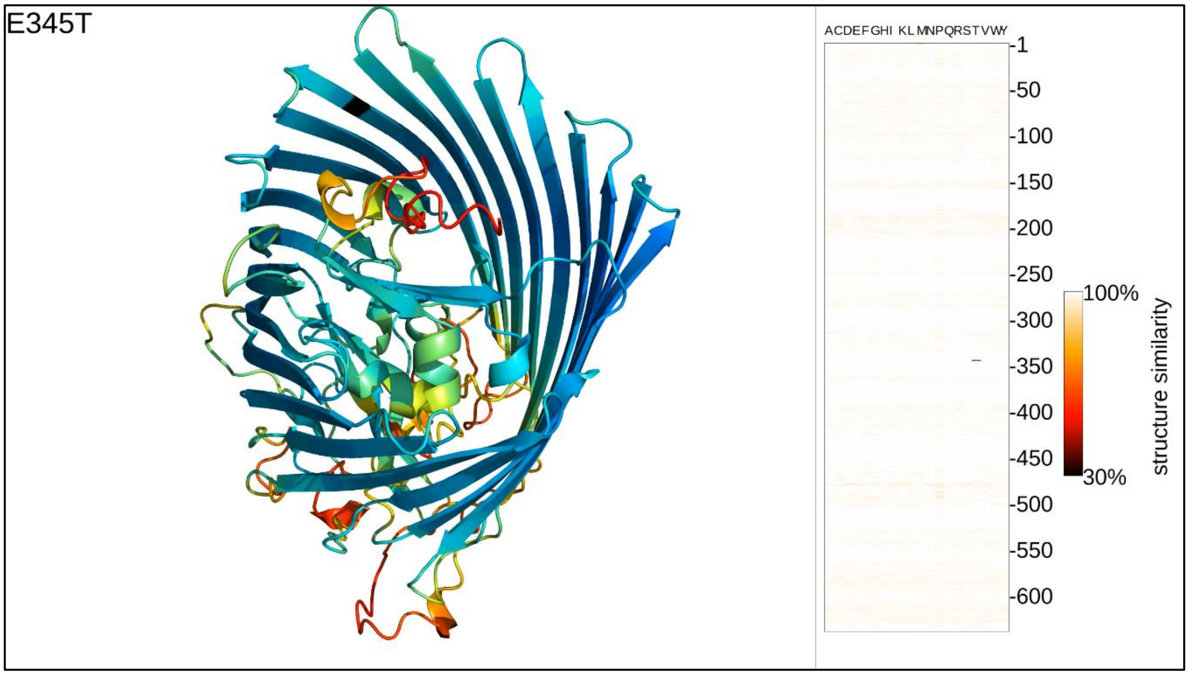

**Fig. S12: Protein mutation movie screenshot: Colicin I receptor Cir from E.coli.** Single frame of a protein mutation movie for Colicin I receptor Cir from E.coli.

#### Table S12: Runtimes for Colicin I receptor Cir from E.coli

| | Runtime (in sec) | |
|---|---|---|
| | 720p resolution | 4K resolution |
| *Prediction (ESMFold)* | 944,335 | |
| *3D rendering* | 8,003 | 33,323 |
| *Structural similarity* | 1,412 | 1,422 |
| *Mutation profile rendering* | 1,129 | 5,217 |
| *Compositing final frames* | 1,183 | 7,731 |
| *Movie rendering* | 35 | 202 |
| *Total (rendering)* | 11,762 | 47,895 |
| *Total (all)* | 956,097 | 992,230 |

**Fig. S13: Prediction runtime increases drastically for longer proteins**

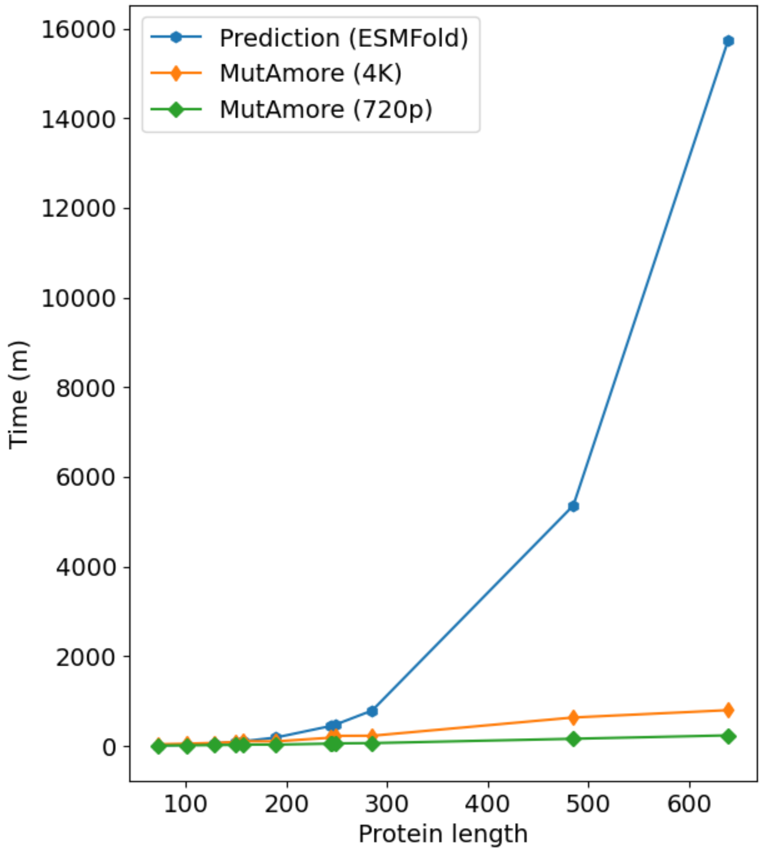

**Fig. S13: Prediction runtime increases drastically for longer proteins.** Mutations for shorter proteins (x-axis; length in residues) can be predicted at high speeds by ESMFold (y-axis; runtime in seconds). For the shortest proteins in the benchmark, MutAmore rendering steps in high resolution (4K) take slightly longer than computing structure predictions. However, above a length of about 200 residues, ESMFold's runtime increases considerably, due to (a) the inability to batch multiple mutant sequences in one computation pass and (b) the general increase in computational complexity and memory consumption of its architecture for longer sequences.
